# High-resolution bulked segregant analysis enables candidate gene identification for bacterial wilt resistance in Italian ryegrass

**DOI:** 10.1101/2023.12.12.571223

**Authors:** Goettelmann Florian, Chen Yutang, Knorst Verena, Yates Steven, Copetti Dario, Studer Bruno, Kölliker Roland

## Abstract

Bacterial wilt, caused by *Xanthomonas translucens* pv. *graminis* (*Xtg*), is a serious disease of economically important forage grasses, including Italian ryegrass (*Lolium multiflorum* Lam.). A major QTL for resistance to *Xtg* was previously identified, but the precise location as well as the genetic factors underlying the resistance are yet to be determined. To this end, we applied a bulked segregant analysis (BSA) approach, using whole-genome deep sequencing of pools of the most resistant and most susceptible individuals of a large (n = 7,484) biparental F_2_ population segregating for resistance to *Xtg*. Using chromosome-level genome sequences as references, we were able to define a ∼300 kb region highly associated to resistance on pseudo-chromosome 4. Further investigation of this region revealed multiple genes with a known role in disease resistance, including genes encoding for Pik2-like disease resistance proteins, cysteine-rich kinases, and RGA4- and RGA5-like disease resistance proteins. Investigation of allele frequencies in the pools and comparative genome analysis in the grandparents of the F_2_ population revealed that some of these genes contain variants with allele frequencies that correspond to the expected heterozygosity in the resistant grandparent. This study emphasizes the efficacy combining BSA studies in very large populations with whole genome deep sequencing and high-quality genome sequences to pinpoint regions associated with a binary trait of interest and accurately define a small set of candidate genes. Furthermore, markers identified in this region hold significant potential for marker-assisted breeding strategies to breed resistance to *Xtg* in Italian ryegrass cultivars more efficiently.

## Introduction

Italian ryegrass (*Lolium multiflorum* Lam.) is a widely cultivated forage species, known for its high yield and nutritional value as well as its fast ground cover (Frame, 1991; Humphreys et al., 2010). As such, it provides an important feed source, as hay or silage, for ruminant livestock production in temperate climates. However, it can be affected by diseases, the most important being bacterial wilt caused by *Xanthomonas translucens* pv. *graminis* (*Xtg*, Egli et al., 1975). The bacteria mainly penetrate through wounds of the plant and spread in the xylem vessels, disrupting the flow of water and nutrients (Rudolph, 1993). This results in wilting and yellowing of the leaves, stunted growth and eventually the death of the plant, causing serious yield losses. Such losses are reported to reach up to 40% under unfavorable conditions (Schmidt and Nuesch, 1980; Gondran and Betin, 1989). Moreover, the global impact of the disease may become more severe due to climate change, which could increase the occurrence of conditions that are favorable to the pathogen, expand its geographical range, or impact host-pathogen interactions (Hunjan and Lore, 2020).

Currently, the only efficient, durable, and economically feasible way to manage the disease is to breed for resistant cultivars. So far, breeding efforts to develop *Xtg* resistant cultivars have been based on recurrent phenotypic selection of resistant individuals following infection with *Xtg*. This has led to cultivars with an increased resistance to the disease (Suter et al., 2021). However, due to the allogamous nature of ryegrasses, these cultivars are genetically heterogeneous, and classical breeding approaches based on phenotypic observation fail to fix dominant alleles, as they can mask susceptibility alleles which are thus maintained in the population. For example, in recurrent selection of *Festuca pratensis*, some level of resistance to *Xtg* was achieved, but quickly stagnated, and complete resistance could not be attained (Boller et al., 2001; Michel, 2001). In this objective, genomics-assisted breeding would greatly aid in identifying and incorporating resistance alleles into elite Italian ryegrass cultivars by targeting specific genetic markers associated with resistance, making the breeding process more efficient. However, such methods require a thorough understanding of the disease resistance and its genetic control.

With the rapid progress in high throughput sequencing technologies in recent years, determining which genetic loci govern a specific phenotype has become increasingly effective and precise. However, such studies, like genome-wide association studies, commonly rely on obtaining both the genotype and phenotype of a large sample of individuals, ranging from hundreds to thousands, which often results in very high costs and efforts (Uffelmann et al., 2021). To reduce costs, bulked segregant analysis (BSA) can be used to identify molecular markers associated with a phenotype of interest in a biparental population using pools of the most extreme phenotypes. The method was first proposed in 1991 and has since undergone significant development in particular thanks to the rise of next generation sequencing (NGS), which allows for high marker densities at a low cost (Michelmore et al., 1991; Li and Xu, 2022).

To statistically determine genetic loci linked to a phenotype of interest, BSA studies generally rely on the ΔSNP index or the G statistics (Magwene et al., 2011; Takagi et al., 2013). However, sequencing data introduces noise associated with many factors, including sequencing errors, varying sequencing coverage or uneven marker density. To reduce this noise, many downstream analyses have focused on developing smoothing methods based on sliding windows. The tricube ΔSNP (t-ΔSNP) index and the G’ statistics, smoothed versions of the ΔSNP index and G statistics, respectively, are based on a weighted average computed by the Nadaraya-Watson tricube smoothing kernel across a sliding window of a set base pair length (Nadaraya, 1964; Watson, 1964). Furthermore, Euclidean distance (ED)-based methods, as well as their smoothed version, ED^4^, have proven to be very powerful at detecting QTL, even outperforming the previous methods (de la Fuente Cantó and Vigouroux, 2022). The implementation of these statistical methods in BSA studies allows to reliably and precisely identify genetic loci associated with a phenotype of interest.

To identify sources of resistance to *Xtg* in *L. multiflorum*, a first study used a segregating F_1_ population (hereafter referred to as Xtg-ART) resulting from a cross between an individual of the susceptible cultivar ‘Adret’ and a resistant individual, M2289, from advanced breeding germplasm (Studer et al., 2006). This study was the first to identify a major QTL explaining 43 to 84% of phenotypic variation for *Xtg* resistance on linkage group (LG) 4. However, further characterization of the QTL was hindered by the predominantly anonymous markers and the lack of genome sequence information for *L. multiflorum* at the time. Later, the assembly of a draft genome sequence of M2289 allowed to apply a BSA approach using shotgun sequencing of pools of the 44 most resistant and the 44 most susceptible individuals within the Xtg-ART population (Knorst et al., 2018a, 2018b). This study was able to confirm the presence of the QTL on LG 4 and highlighted the efficacy of BSA in fine-mapping the QTL for resistance. However, the small number of individuals available, together with the low depth of the sequencing output and the fragmented nature of the M2289 draft genome sequence prevented further fine-mapping of the QTL region and determination of the underlying genes. Recently the high-quality chromosome-level genome assembly of the cultivar ‘Rabiosa’ for *L. multiflorum* was made available, providing a strong basis for further genomics-based analyses in Italian ryegrass (https://www.ncbi.nlm.nih.gov/datasets/genome/GCA_030979885.1/). Furthermore, the high-throughput sequencing platforms currently available allow for a high coverage of the whole genome sequence, resulting in high marker densities and more accurate allele frequency estimations, improving the effectiveness and accuracy of BSA studies.

The goal of this study was to define candidate genes for resistance to *Xtg* by applying a BSA approach based on high coverage whole-genome sequencing of pools of susceptible and resistant individuals in an F_2_ population derived from the Xtg-ART population, hereafter referred to as the Xtg-ART-F_2_ population. We aimed at fine-mapping the major QTL on LG 4, identifying candidate resistance genes, and defining markers associated with resistance to be used in marker-assisted breeding.

## Results

### Resistance to bacterial wilt is segregating in the Xtg-ART-F_2_ population

The disease progress in the 7,484 Xtg-ART-F_2_ individuals in the greenhouse followed the expectations, with first symptoms being visible at 14 days post infection (dpi) and becoming more severe over time (Fig. 1). Since the natural death of older leaves was challenging to distinguish from *Xtg* symptoms, most healthy plants were assigned a score of two at 21 and 28 dpi. Cutting back the plants after the third scoring allowed to make this distinction between natural leaf senescence and *Xtg* symptoms and at 49 dpi 1,165 (15.5%) plants showed no symptoms, while 1,140 (15.2%) had died. From these two sets of plants, the most resistant and the most susceptible individuals were selected, resulting in a total of 750 resistant plants and 761 susceptible plants that were used to form two pools that were sequenced.

**Figure 1.**
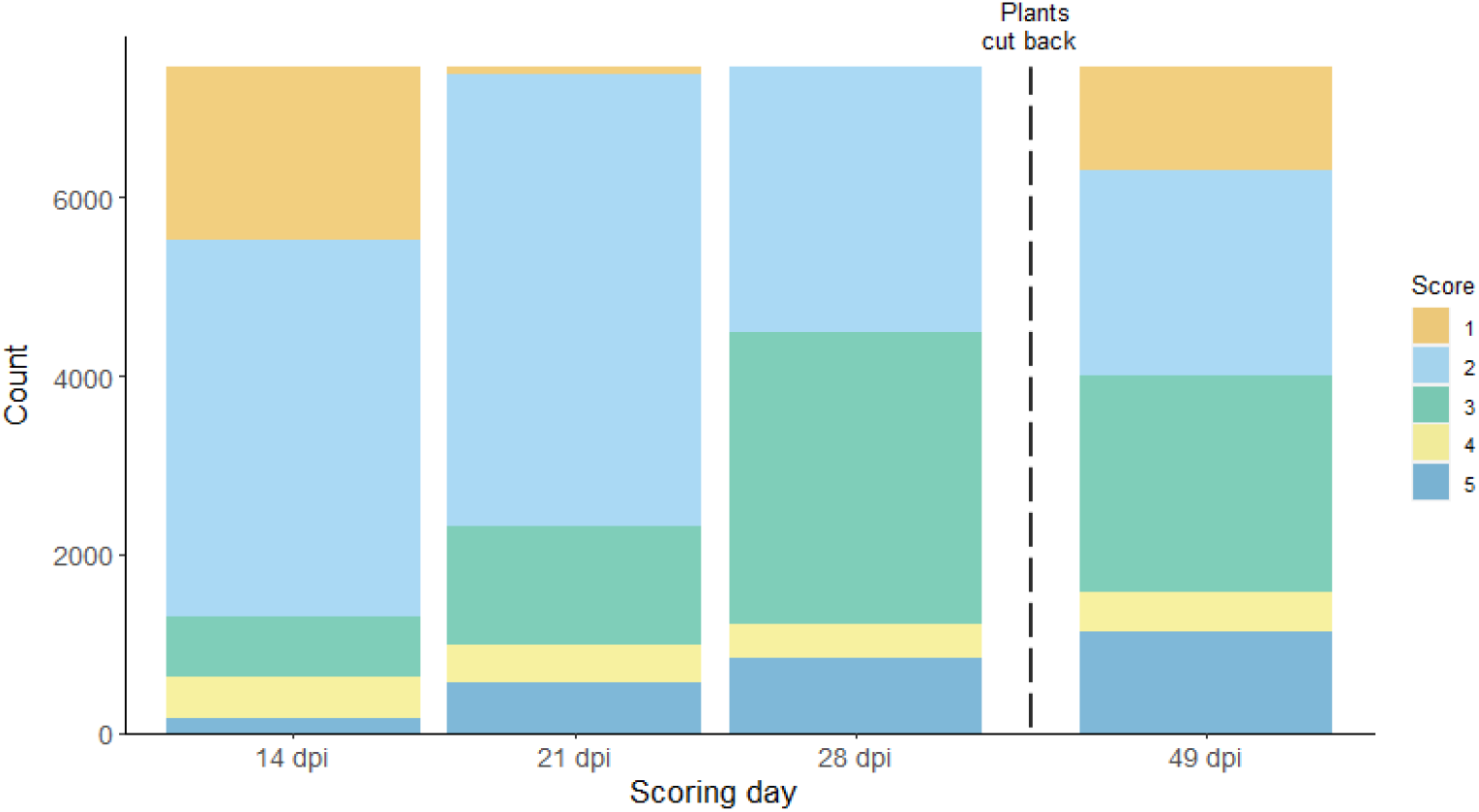
Distribution of the bacterial wilt disease scores in 7,484 individuals of the *L. multiflorum* Xtg-ART-F_2_ population at 14, 21 and 28 days post inoculation with *Xtg* (dpi), as well as at 49 dpi, three weeks after the plants were cut back and allowed to regrow. A score of 1 corresponds to a healthy plant showing no symptoms (orange), 2 corresponds to mild symptoms (light blue), 3 corresponds to intermediate symptoms (green), 4 corresponds to severe symptoms (yellow), and 5 corresponds to a dead plant (dark blue).

### Obtaining chromosome-level genome sequences of the Xtg-ART-F_2_ grandparents

In order to obtain a chromosome-level genome sequence that is representative of the resistant individuals of the Xtg-ART-F_2_ population, the draft genome assembly of the resistant grandparent M2289 (Knorst et al., 2018b), which consists of 129,579 scaffolds, was anchored to the chromosome-level Rabiosa genome sequence (https://www.ncbi.nlm.nih.gov/datasets/genome/GCA_030979885.1/). A total of 78,219 scaffolds were successfully anchored to the seven pseudo-chromosomes of the Rabiosa assembly, and 3,912 scaffolds from the M2289 assembly were anchored to 829 larger scaffolds that are not anchored to pseudo-chromosomes in the Rabiosa assembly. In addition, 47,448 scaffolds from the M2289 assembly could not be anchored and were excluded from further analysis. This chromosome-level genome sequence of M2289 will hereafter be referred to as “M2289_v2”.

To compare M2289_v2 with the genome sequence of the susceptible grandparent of the Xtg-ART-F_2_ population, a similar approach was applied to the Adret draft genome assembly, which consists of 117,277 scaffolds, to produce the “Adret_v2” genome sequence (Knorst et al., 2018a). A total of 72,888 scaffolds were anchored to pseudo-chromosomes, 3,596 scaffolds were anchored to 795 larger scaffolds from the Rabiosa assembly, and 40,793 scaffolds were left untouched.

### A 300 kb region of pseudo-chromosome 4 is highly associated with resistance

A total of 606,760 Gb of DNA sequence information was obtained by whole-genome sequencing of the two pools (average = 303,378 Gb per pool), corresponding to an average of 120× coverage of the haploid genome per pool, considering an estimated genome size of 2.5 Gb (Copetti et al., 2021). After trimming and filtering reads, a total of 490,386 Gb remained (average = 245,193 Gb per pool), corresponding to an average of 98.1× coverage of the haploid genome per pool. The mapping of the reads to the M2289_v2 genome sequence resulted in an average alignment rate of 34.13%. The variant calling yielded a total of 7,965,619 biallelic variants, including 7,068,640 SNPs and 896,979 INDELs. After filtering for total depth and quality, a total of 4,239,024 variants remained for the subsequent analysis. Finally, to have a comparison with a better quality and more complete genome sequence, the Rabiosa genome sequence was also used as a reference, which resulted in an average of 87.77% alignment rate, yielding a total of 18,765,678 biallelic variants, including 15,170,193 SNPs and 3,595,485 INDELs. After filtering, 7,525,443 variants remained for the subsequent analysis.

All three approaches used to test the association of each variant to resistance, the |ΔSNP| index, the G value and the ED value, showed a clear association on pseudo-chromosome 4, with a high peak spanning most of the pseudo-chromosome (Figure 2). However, a high level of noise can be seen, with fluctuations attributable to sequencing errors or the stochastic nature of sequencing. This results in many interspersed regions with high association to resistance, that are unexpected in a biparental population, where a broad peak of association to resistance is expected from a segregation distortion. Consequently, these fluctuations hindered the precise identification of the region where association to resistance was the highest on pseudo-chromosome 4.

**Figure 2.**
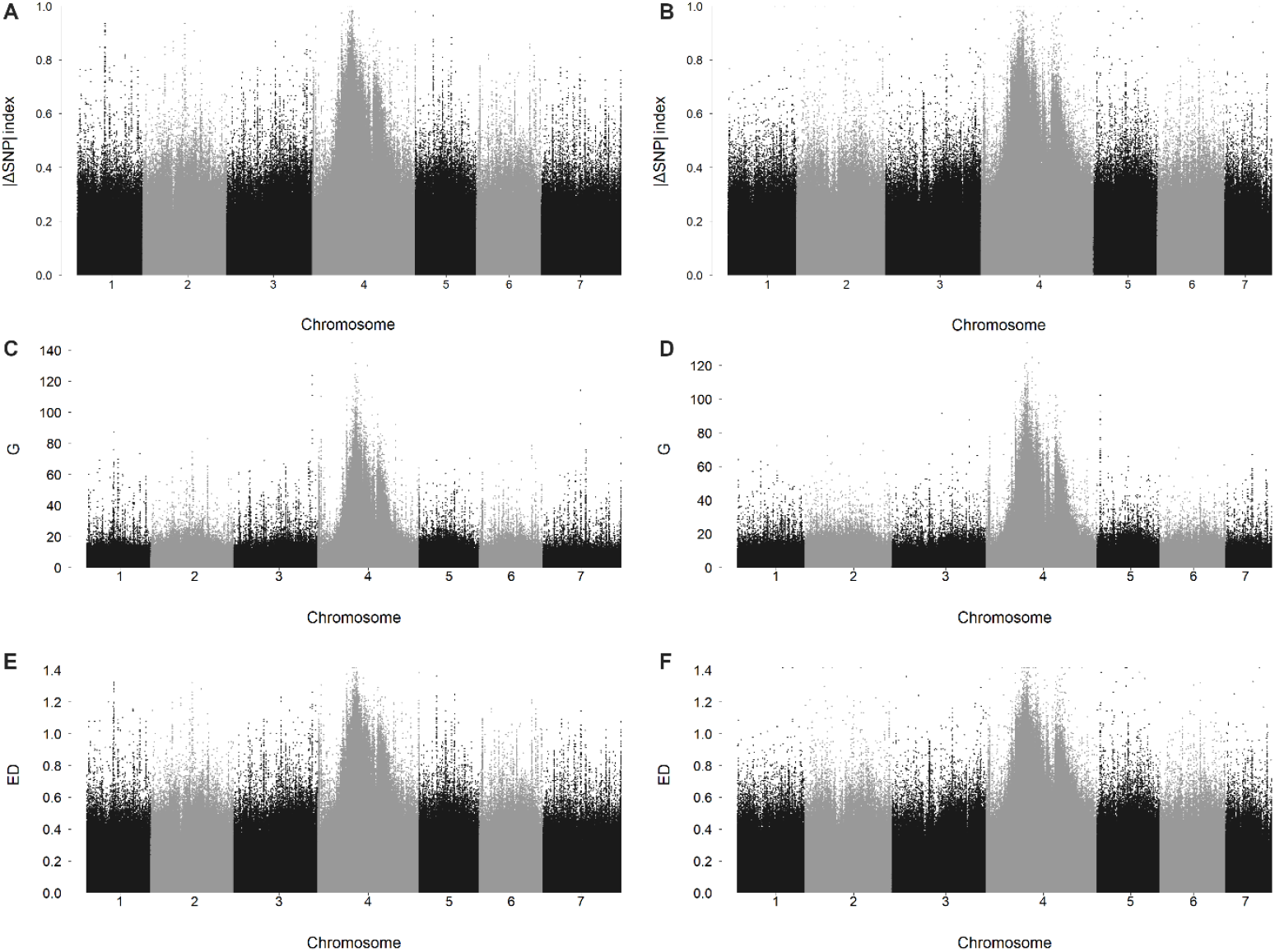
Association of each variant to *Xanthomonas translucens* pv. *graminis* resistance across all seven *Lolium multiflorum* pseudo-chromosomes calculated with the |ΔSNP| index (A, B), the G value (C, D) or the Euclidean Distance (ED) (E, F) with M2289_v2 (A, C, E) or Rabiosa (B, D, F) as a reference genome sequence.

To increase the signal to noise ratio as well as to refine the peak region, smoothing methods based on sliding windows were applied to obtain the t-|ΔSNP| index, the G’ value and the ED^4^ value. These methods effectively reduced noise and allowed to distinguish a clear peak on pseudo-chromosome 4, which exceeded the 99% confidence interval for the t-|ΔSNP| index and the 99% false discovery rate threshold for the G’ value (Figure 3). With all approaches, the strongest association to resistance was found at a similar location on pseudo-chromosome 4, and no association to resistance was found on any other pseudo-chromosome. With the ED^4^ approach, a sharp peak could be seen, spanning approximately from 28,800,000 bp to 28,900,000 bp in the M2289_v2 genome sequence, and this region was selected to be further investigated (Figure 3 E, F). In Rabiosa, similar results were observed, and a sharp peak was observed from approximately 163,280,000 bp to 163,400,000 bp with the ED^4^ approach (Suppl. Figure 1).

**Figure 3.**
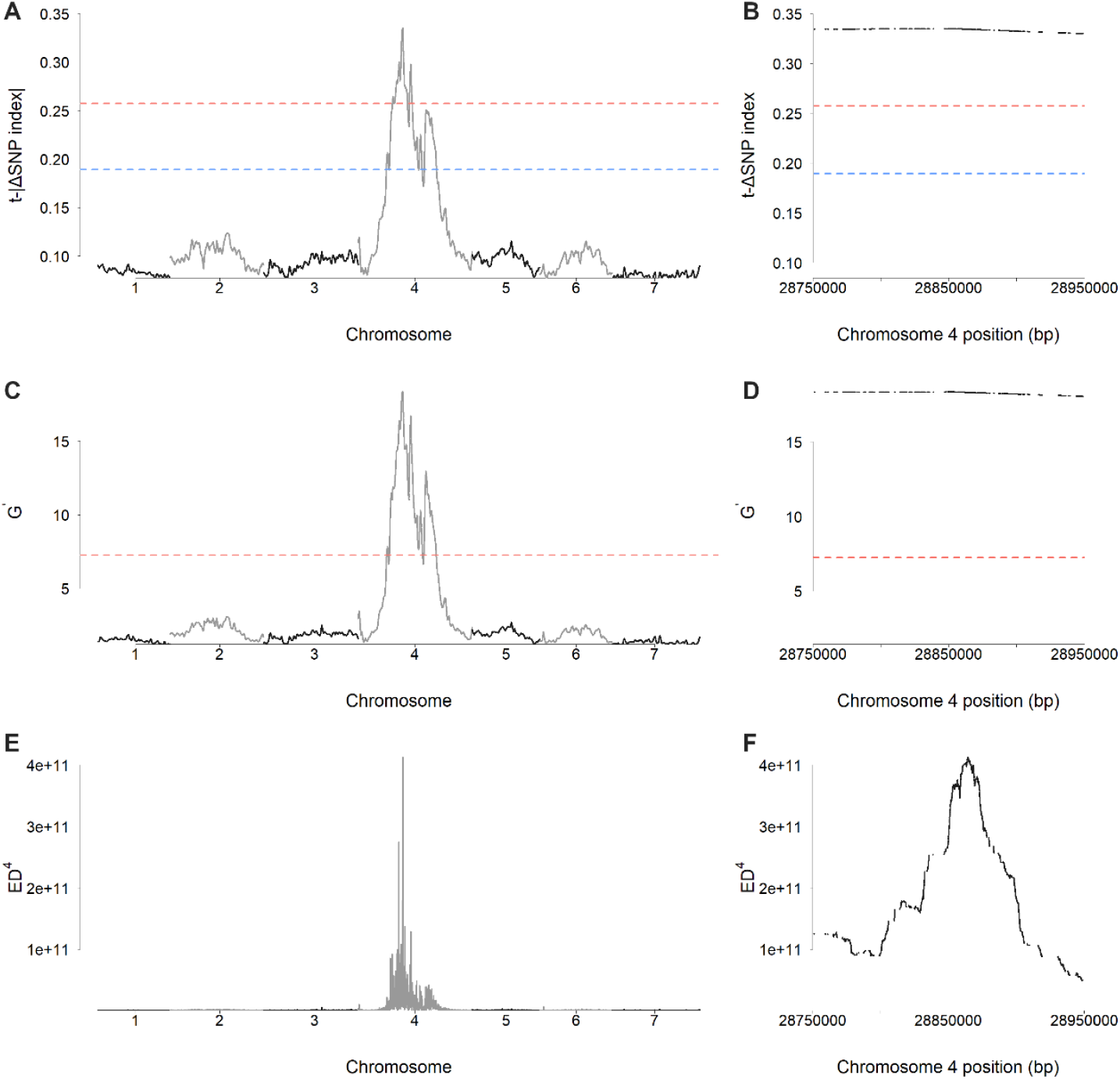
Association of each variant to *Xanthomonas translucens* pv. *graminis* resistance in *Lolium multiflorum* calculated with the t-|ΔSNP| index (A, B), the G’ value (C, D) and the ED^4^ value (E, F), with M2289_v2 as a reference genome sequence, on all pseudo-chromosomes (A, C, E) or the region with the highest association to resistance on pseudo-chromosome 4 (B, D, F). Dotted blue lines represent the 95% confidence interval for the t-|ΔSNP| index, and dotted red lines represent the 99% confidence interval for the t-|ΔSNP| index or the false discovery rate threshold of 1% for the G’ value.

A BLASTn-based comparison of the annotated gene sequences within these two regions was able to determine that the region identified in Rabiosa corresponded to a portion of the region identified in M2289_v2. For a better comparability of the two regions, the larger region in Rabiosa, corresponding to the M2289_v2 region, was defined, spanning from 163,094,000 bp to 163,410,000 bp and selected for further analysis. These regions were investigated in more detail and revealed a cluster of genes that could potentially be linked to disease resistance.

### Multiple candidate resistance genes were found in the candidate region

A total of twelve and nine annotated genes, found in the candidate regions of M2289_v2 and Rabiosa, respectively, showed similarity to genes involved in disease resistance, and were defined as candidate genes for resistance to *Xtg* (Table 1). These included genes showing similarity to Pik-2-like and resistance gene analog (RGA)4-like disease resistance proteins, as well as two cysteine-rich kinases (CRK) that were found in both genome sequences. The candidate region in the M2289_v2 assembly also contained two RGA5-like disease resistance proteins, while in the Rabiosa assembly, four serine protease inhibitor (SERPIN) genes were found. Additionally, in Rabiosa, one gene showed homology to a retrotransposon, and one to a histone deacetylase, which could not be linked to disease resistance. Furthermore, eight and four genes, in M2289_v2 and Rabiosa respectively, could not be annotated as no match to any known gene was found. Moreover, in M2289_v2, the region contained 17 gaps and spanned ∼100 kb, whereas the corresponding region in the Rabiosa assembly spanned ∼300 kb without any gaps. This indicates that additional genes could be present in this region but cannot be found in the M2289_v2 assembly.

**Table 1:** Genes found within the candidate region, defined from 28,800,000 bp to 28,900,000 bp on pseudo-chromosome 4 of the M2289_v2 genome sequence, and from 163,090,000 bp to 163,410,000 bp on pseudo-chromosome 4 of the Rabiosa genome sequence, that were annotated as a potential disease resistance gene.

| Annotation | M2289_v2 gene ID | Start (bp) | End (bp) | Rabiosa gene ID | Start (bp) | End (bp) |
| --- | --- | --- | --- | --- | --- | --- |
| Pik-2-like | XLOC_040221 | 28,810,090 | 28,810,476 | evm.TU.chr4.21718 | 163,094,711 | 163,100,016 |
|  | XLOC_040220 | 28,811,063 | 28,811,263 |  |  |  |
|  | XLOC_040219 | 28,811,461 | 28,811,664 |  |  |  |
|  | XLOC_040218 | 28,812,525 | 28,812,767 |  |  |  |
|  | XLOC_000927 | 28,814,543 | 28,815,826 |  |  |  |
|  | XLOC_000926 | 28,816,808 | 28,816,882 |  |  |  |
|  | XLOC_021628 | 28,818,511 | 28,819,200 |  |  |  |
| CRK6 | XLOC_017025 | 28,830,345 | 28,831,876 | evm.TU.chr4.21727 | 163,153,306 | 163,154,874 |
| CRK8 | XLOC_023208 | 28,853,351 | 28,854,622 | evm.TU.chr4.21737 | 163,276,545 | 163,286,601 |
|  |  |  |  | evm.TU.chr4.21738 | 163,325,147 | 163,333,991 |
| SERPIN |  |  |  | evm.TU.chr4.21747 | 163,365,239 | 163,366,661 |
|  |  |  |  | evm.TU.chr4.21748 | 163,368,491 | 163,369,111 |
|  |  |  |  | evm.TU.chr4.21749 | 163,369,308 | 163,385,542 |
|  |  |  |  | evm.TU.chr4.21750 | 163,385,641 | 163,387,209 |
| RGA5-like | XLOC_055978 | 28,892,028 | 28,892,210 |  |  |  |
|  | XLOC_055976 | 28,895,207 | 28,895,509 |  |  |  |
| RGA4-like | XLOC_055977 | 28,899,358 | 28,901,523 | evm.TU.chr4.21754 | 163,406,159 | 163,409,767 |

The Pik-2-like disease resistance protein gene evm.TU.chr4.21718 from Rabiosa was homologous to the region in M2289_v2 where seven Pik-2-like genes were found, indicating that they may correspond to a single gene, homologous to the one found in Rabiosa (Figure 4). Additionally, regions displaying homology to the two genes that showed similarity to RGA5-like disease resistance proteins in M2289_v2 were found in Rabiosa, although they were not predicted as genes in the Rabiosa annotation (Figure 4). Overall, the comparison of these candidate genes between M2289_v2 and Rabiosa, as well as with the updated genome sequence of the susceptible grandparent, Adret_v2, showed that most genes are conserved between the three genome sequences. Still, no homology to the evm.TU.chr4.21738 and evm.TU.chr4.21749 genes from Rabiosa was found in M2289_v2 nor Adret_v2, and homology to evm.TU.chr4.21750 was found only in Adret_v2.

**Figure 4.**
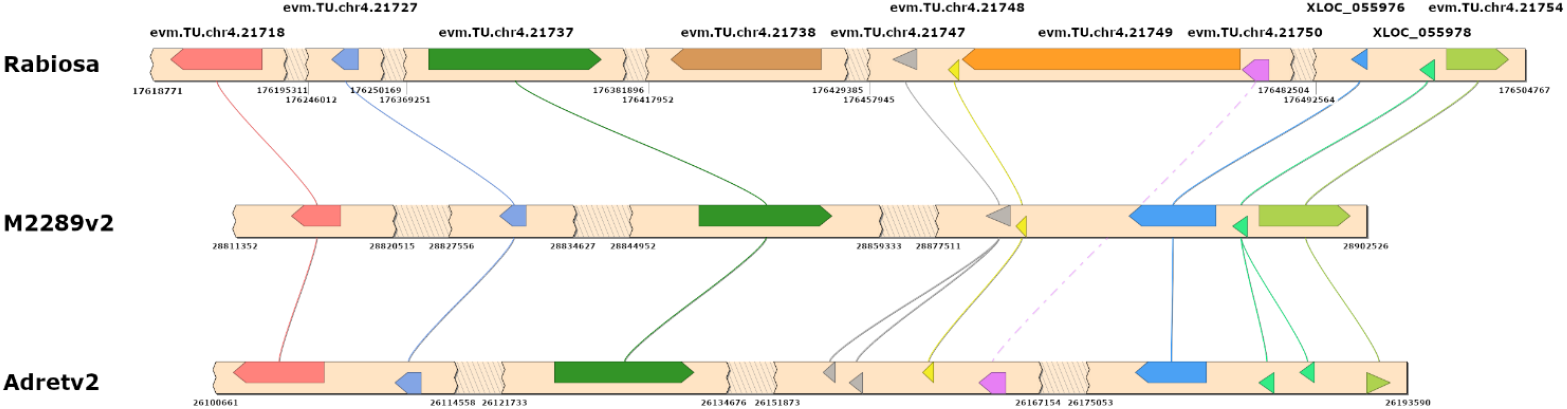
Comparison of the candidate genes composition within the candidate region across the genome sequences of Rabiosa, the resistant grandparent M2289_v2, and the susceptible grandparent Adret_v2. Genes are represented as bars, with the arrowhead representing the 3’ end of the gene. Synteny between genome sequences is displayed as lines connecting homologous genes. Regions with no genes are shown as hatched breaks for improved clarity.

A total of 3,213 variants were found in the candidate region using M2289_v2 as a reference, and 3,177 with Rabiosa. Of these, 880 and 1,010, respectively, had a variant that was absent or present in very low frequency in the susceptible pool and the Adret susceptible grandparent, corresponding to an absence or low frequency expected of a dominant resistance allele in a susceptible individual or population. Using M2289_v2 as a reference, 101 of these variants were found in coding sequences of 11 genes, and 54 were found to induce a nonsynonymous amino acid change, in a total of eight genes (Figure 5A). The allele frequencies in XLOC_000927, XLOC_021628 and XLOC_017025 indicate that these three genes are heterozygous at these loci in the M2289 grandparent. The remaining five genes, however, seem to be homozygous in M2289. With the Rabiosa assembly, 74 variants with a resistance allele were found in a coding sequence, in a total of six genes, and 47 would result in a nonsynonymous amino acid change, found in the same six genes (Figure 5B). Most of these variants were homozygous in the M2289 resistant grandparent, with only eight being heterozygous in evm.TU.chr421718, evm.TU.chr421747 and evm.TU.chr421750.

**Figure 5.**
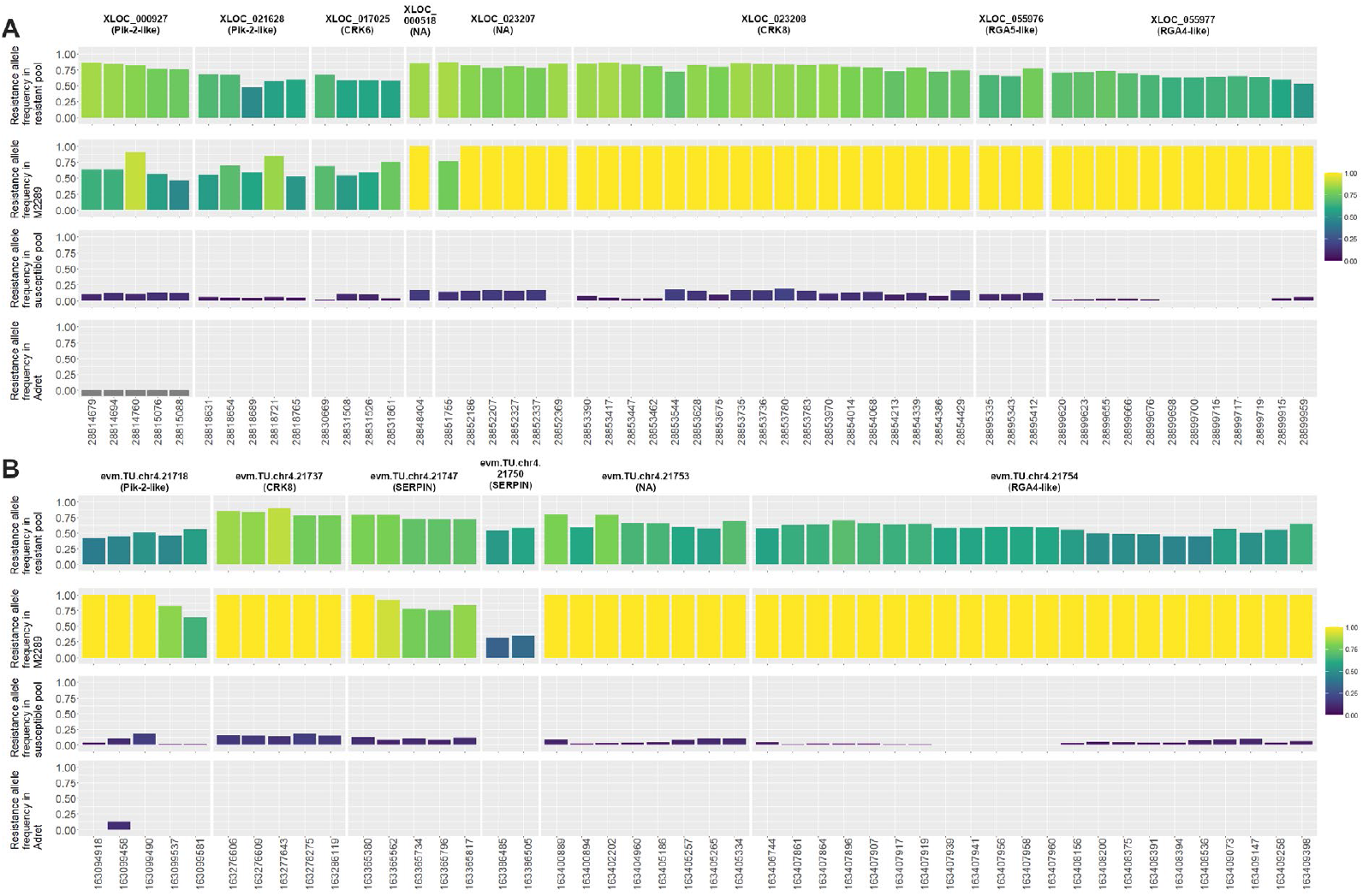
Frequencies of resistance alleles for variants causing amino acid changes in annotated genes within the candidate region. Frequencies are represented within the resistant and susceptible pools, as well as within the M2289 and Adret grandparents, with both the M2289_v2 (A) and the Rabiosa (B) genome sequences. Gray negative bars denote an absence of reads for this variant.

## Discussion

We were able to define a ∼300 kb genome region that was highly associated with resistance to bacterial wilt in *L. multiflorum*. This region contained a small number of candidate resistance genes with known roles in disease resistance. Consistent results were found with three different statistics for BSA data analysis, supporting the robustness of the applied methodology. It also demonstrates the efficacy of BSA based on whole genome sequencing of pools of contrasting phenotypes to identify candidate genes associated with a binary trait of interest in a *L. multiflorum* biparental F_2_ population.

While previous work based on a first draft of the M2289 genome sequence was able to identify genomic regions associated with resistance mapping to chromosome 4, the very fragmented nature of this genome assembly, as well as the lack of reliable information on the position of each scaffold on chromosomes, did not allow for a precise location of the resistance gene in the genome (Knorst et al., 2018a; Goettelmann et al., 2021). The recent development of a high-quality chromosome-level genome sequence of Rabiosa allowed to produce the updated chromosome-level genome sequence M2289_v2. Though this genome sequence is still fragmented, it allowed to reliably anchor scaffolds on a pseudochromosome, which enabled a clear visualization of the association to bacterial wilt resistance across the genome and allowed to narrow down the location of the QTL on pseudo-chromosome 4. Furthermore, the sliding window-based statistical approaches used in this study required highly contiguous genome sequences and could not have been applied with the M2289 draft genome sequence, in which the largest scaffold is 43,903 bp, much smaller than the window size of one million bp we used (Knorst et al., 2018b).

The high population size coupled with a whole-genome sequencing approach at high coverage was able to optimize most of the factors on which the success of BSA studies rely, (Magwene et al., 2011; de la Fuente Cantó and Vigouroux, 2022; Huang et al., 2022). In BSA studies that have been performed so far, the sizes of the investigated populations were generally small, often around 200 individuals, though they can range from <100 to >10,000 individuals (reviewed in Huang et al., 2022). Pool sizes were also variable, ranging from only five individuals per pool to more than 400. In our study, the investigation of a total of 7,484 F_2_ individuals, and the selection of 750 and 761 individuals for the resistant and susceptible pool, respectively, is thus relatively high and allowed for both a high detection power and mapping resolution. Another crucial factor for BSA studies is the number of recombination events. This is of particular importance as *Lolium* species have been shown to exhibit a low recombination rate, in particular in chromosome 4 (Kopecký et al., 2010; King et al., 2013). Nonetheless, the use of an F_2_ population and the high number of individuals per pool allowed to effectively address this factor.

Using both the M2289_v2 and the Rabiosa genome sequences as reference showed consistent results, with the highest association to resistance found in a similar location in both genome sequences. The genes found within the two regions were comparable, both in terms of presence of the genes, as well as their order. Still, some genes found in Rabiosa were not found in M2289_v2. While they may be absent from the M2289 genome, this may also be explained by the fragmented nature of this genome sequence, with many gaps present in the candidate region. These genes may either be missing from the assembly or may not have been correctly anchored when producing the M2289_v2 genome sequence. Furthermore, the Rabiosa genotype shows some susceptibility to *Xtg* and may not contain the resistance source (unpublished data). Therefore, the anchoring of the M2289 genome sequence to Rabiosa may not allow to properly access the resistance source in M2289. In human genetics, where one genome sequence is universally used as a reference, many population-specific DNA sequences were missing from this reference, preventing the identification of genes of interest (Sherman et al., 2019). This highlights the need for high-quality, complete genome assemblies that are related to the investigated population in association studies. Therefore, obtaining the complete DNA sequence of resistant individuals at the candidate region would be crucial to accurately determine genes that could be responsible for resistance to *Xtg*.

Nonetheless, the present study was able to identify multiple genes that are known to be involved in disease resistance. The most interesting candidates within the genome region associated with *Xtg* resistance include XLOC_000927, XLOC_021628 and evm.TU.chr4.21718, annotated as Pik-2-like disease resistance proteins. Pik proteins are nucleotide-binding leucine-rich repeat (NLR) proteins that were shown to be involved in resistance to many strains of rice blast (*Magnaporthe grisea*), recognizing the AVR-Pik proteins from the pathogen and triggering a specific defense response (Ashikawa et al., 2012; Wang et al., 2017). However, the evm.TU.chr4.21718 gene from Rabiosa showed homology to the M2289_v2 region containing seven genes annotated as Pik-2-like disease resistance proteins, suggesting these seven genes may correspond to a single gene in M2289_v2. The prediction of multiple smaller genes at this locus in M2289_v2 is likely due to the fragmented nature of the genome sequence, since this gene region is found on three separate scaffolds in the original draft assembly, from which the annotation was produced. Another candidate is XLOC_017025, annotated as a cysteine-rich receptor-like kinase (CRK) 6. These proteins are transmembrane proteins involved in the detection of biotic and abiotic stresses, leading to programmed cell death (Chen et al., 2004). In *A. thaliana*, the overexpression of *CRK6* and other *CRK* genes led to an enhanced activation of pathogen-associated molecular pattern-triggered immunity and some CRK have been shown to mediate pattern-triggered immunity (Ederli et al., 2011; Yeh et al., 2015). Finally, evm.TU.chr4.21747 and evm.TU.chr4.21750, two candidate genes of the serine protease inhibitor (SERPIN) superfamily, were found. SERPINs act on many cell processes, and some members of the family have been shown to have a role in disease resistance in *A. thaliana* and barley (*Hordeum vulgare* L.) by inhibiting proteinases from the pathogen (Pekkarinen et al., 2007; Laluk and Mengiste, 2011; Quilis et al., 2014). Moreover, while the allele frequencies of variants found in these genes were not in line with the expected heterozygous frequency in the M2289 parent, the candidate region also contained genes encoding for CRK8, a member of the CRK family like CRK6. Additionally, two other SERPINs in Rabiosa, that could be involved in resistance were also found. Lastly, the candidate region also contained genes of the RGA4- and RGA5-like protein families. These NLR proteins act together in mediating disease resistance to *M. grisea* in rice (*Oryza sativa* L.) Césari et al., 2014).

The identified candidate region provides valuable insight into the potential loci responsible for resistance to *Xtg*. Interestingly, this region contains mostly genes with known functions in disease resistance. As such genes are often found in clusters containing other resistance genes as well as paralogs resulting from the duplication and divergence of some genes, it is plausible that multiple genes within this region could play a role in resistance to *Xtg* (Michelmore and Meyers, 1998; Sekhwal et al., 2015). This emphasizes the need for a more comprehensive overview of the genes present in the candidate region in resistant individuals to better understand the mechanisms involved in resistance to *Xtg*.

Ultimately, this study allowed to identify genetic variants that were highly associated with resistance to *Xtg* and could be used as diagnostic markers for marker-assisted selection. In wheat (*Triticum aestivum* L.), markers associated with the *Fhb1* QTL for resistance to *Fusarium* head blight are routinely used in breeding programs, significantly improving resistance to the disease in new cultivars (Anderson et al., 2007). In rice, marker-assisted selection allowed the introgression of the *Xa38, Xa21* and *xa13* resistance genes against the bacterial blight pathogen *Xanthomonas oryzae* pv. *oryzae*, as well as the *Pi2* and *Pi54* resistance genes against rice blast caused by *M. oryzae*, into elite rice cultivars (Swathi et al., 2019; Yugander et al., 2019; Sagar et al., 2020). Although further validation is necessary to confirm their effectiveness, the identified variants hold significant potential as molecular markers to be used in marker-assisted breeding applications, laying the foundations for an efficient selection of bacterial wilt resistance in Italian ryegrass cultivars.

### Experimental procedures

#### Plant material, bacterial material, inoculation

A total of 143 F_1_ individuals from the Xtg-ART population (Studer et al., 2006) were planted in isolation tents for a polycross in late summer of 2017 and an average of 6,000 seeds were harvested from each F1 plant in early summer of 2018. Self-pollination was identified in the F1 plants using a fully informative single sequence repeat (SSR) marker, and 131 F1 plants were confirmed to come from cross-pollination and their seeds were selected for the experiment. An equal number of 61 seeds per F1 plant (7,991 seeds in total) were planted in 96 30×45 mm pot trays filled with soil and grown in the greenhouse. From these, 7,484 individual plants were established and formed the Xtg-ART-F_2_ population. After a month, plant material was harvested from each F_2_ plant and stored individually at - 80°C. The plants were allowed to grow back for a month before being inoculated.

For the inoculation, bacteria from the highly virulent Xtg29 strain (Kölliker et al., 2006) were grown at 28°C on YDC agar medium (2% dextrose, 1% yeast extract, 2% CaCO_3_, 1.5% agar) for 48 h. Bacteria were then dissolved in a physiological solution (0.877% NaCl) and the OD_580_ was adjusted to 0.3, corresponding to a concentration of 10^8^ CFU/mL.

The plants were inoculated by dipping scissors in the bacterial solution and cutting them 4 cm above soil. Each plant was then scored at 14, 21 and 28 days post infection (dpi). The plants were then cut back and allowed to grow back, and a last scoring was performed at 49 dpi. Scorings were performed on a scale of one to five, where one corresponds to a healthy plant showing no symptoms and five corresponds to a dead plant.

At the last scoring, 1,165 plants showed no symptoms and 1,140 were dead (Figure 1). From these, to select truly resistant and susceptible plants, inconsistent scores across the scoring period were filtered out. For the resistant individuals, plants showing a score of three or above anytime during the scoring period were excluded, and a total of 750 plants remained. For the susceptible individuals, plants showing a disease score of three and above at early stages (14 and 21 dpi) and of four and above at a later stage (28 dpi) were selected, accounting for a total of 761 plants.

### DNA extraction and Sequencing

DNA was extracted from the previously harvested plant material using the NucleoSpin® Plant II DNA kit (Macherey Nagel, Duren, Germany). One leaf cutting of approximately three centimeters per plant was used. For a better quality and accuracy of the sequencing, each pool was divided into three sequencing libraries of an equal number of individuals (∼250) that were sequenced independently and pooled after sequencing. Sequencing libraries were prepared using the Illumina TruSeq Nano DNA Library Preparation, (Illumina, San Francisco, CA, USA) and sequenced using 150 bp paired-end reads on one lane of an S4 flowcell using the Illumina NovaSeq 6000 platform by the Functional Genomics Center Zurich, Zurich, Switzerland.

### Production of chromosome-level genome sequences, alignment and variant calling

A chromosome-level version of the M2289 resistant grandparent, M2289_v2, was produced by anchoring the scaffolds of the published M2289 draft genome sequence to the recently-obtained high-quality assembly of the *L. multiflorum* cultivar ‘Rabiosa’ (https://www.ncbi.nlm.nih.gov/datasets/genome/GCA_030979885.1/) using RagTag ‘scaffold’ v2.1.0 with default parameters (Knorst et al., 2018b; Alonge et al., 2019, 2021). The same approach was used to anchor the draft genome sequence of the Adret susceptible grandparent, resulting in the chromosome-level Adret_v2 genome sequence (Knorst et al., 2018a). Additionally, the subsequent analysis was also applied using the Rabiosa genome sequence as reference.

The obtained reads were trimmed to remove adapter sequences and low-quality bases with trimmomatic v0.35 using TruSeq adapter sequence information and options “LEADING:3 TRAILING:3 SLIDINGWINDOW:4:15 MINLEN:70” (Bolger et al., 2014). The trimmed reads were then aligned to the M2289_v2 or the Rabiosa genome sequence with bowtie2 v2.3.4.2 using parameter “--no-unal” (Langmead and Salzberg, 2012). The resulting alignments were converted into binary alignment files and sorted using samtools v1.9 (Danecek et al., 2021). Variants were identified using samtools ‘mpileup’ and bcftools ‘call’ with option “-t DPR” for depth information (Danecek et al., 2021).

The resulting variant call file was then loaded into R version 4.1.0 with the ‘vcfr’ package v1.12.0 (R Core Team, 2022; Knaus and Grünwald, 2017). Loci with a QUAL score lower than 50 were excluded, and only biallelic loci were retained. Variants were filtered for read numbers >50 and <200 (summed across both pools).

### Identification and characterization of candidate genes

For each variant, the Euclidean distance (ED) between the resistant and the susceptible pool was calculated as defined by Hill et al., 2013, modified for biallelic loci as follows:

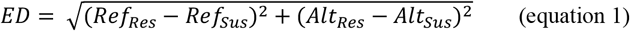

where Ref_Res_ and Ref_Sus_ are the frequencies of the reference allele in the resistant pool and in the susceptible pool, respectively, and Alt_Res_ and Alt_Sus_ are the frequencies of the alternative allele in the resistant pool and in the susceptible pool, respectively. To increase the signal to noise ratio, the ED^4^ was calculated for each variant as the fourth power of the sum of ED within a window of 1,000 consecutive variants, centred around this variant. As the ΔSNP approach requires the reference genome used to be of a parent of the population, we used the absolute value of the ΔSNP (|ΔSNP|) to compare the analysis with M2289_v2 and Rabiosa as reference genomes, as proposed by Omboki et al., 2018. The |ΔSNP| index and the G value, as well as their corresponding smoothed values, the t-|ΔSNP| index and G’, were calculated with the QTLseqR package, using a sliding window size of 10^6^ for the M2289_v2 analysis and 5×10^6^ for the Rabiosa analysis, as this genome sequence is larger and more complete (Mansfeld and Grumet, 2018). The results were then visualized in Manhattan plots with the ‘qqman’ R package (Turner, 2018)

Candidate genes were defined by determining the genes present in the identified regions in the genome annotations of both M2289_v2 and Rabiosa. The regions identified with both reference genome sequences were then compared by performing a BLASTn v. 2.2.30 using the candidate gene sequences as query and the genome sequences as subject. This showed that the peak region in Rabiosa corresponds to a subset of the region identified in M2289_v2. For a better comparison, the larger region in Rabiosa corresponding to the region in M2289_v2 was selected, and the list of genes appended accordingly. The identified genes were then annotated by performing a BLASTx with the gene nucleotide sequences as query and the TAIR10 *A. thaliana* or the NCBI nr protein databases as subject (Berardini et al., 2015; Sayers et al., 2022). Genes that showed homology to genes known to be involved in disease resistance were defined as candidate genes. The synteny between M2289_v2 and Rabiosa was determined with a BLASTn-based approach using the SimpleSynteny online tool (www.dveltri.com/simplesynteny/) with the sequences of candidate genes in Rabiosa, as this assembly is more complete, as well as two additional genes that were identified in M2289_v2 only (Table 1, Veltri et al., 2016).

To further characterize the candidate genes, raw reads from the M2289 and Adret grandparents of the were aligned to the M2289_v2 and the Rabiosa reference genome sequences and variants were called as before (Knorst et al., 2018a, 2018b). Candidate genes containing variants were defined and the allele frequencies of the variants investigated. As it is expected that the resistance allele is absent in susceptible individuals, loci where one allele had a frequency < 0.2 in the susceptible pools and the Adret susceptible grandparent were identified, and this allele was then defined as a resistance allele. Loci where such variants would result in a change in amino acid were then determined. The frequency of resistance alleles in these loci was then investigated in both pools, M2289 and Adret.

## Data availability

The *Lolium multiflorum* genome ‘Rabiosa’ is available on https://www.ncbi.nlm.nih.gov/datasets/genome/GCA_030979885.1/. The chromosome-level genome sequences of the Xtg-ART-F_2_ grandparents (M2289_v2 and Adrev2) generated in this study are available on https://doi.org/10.5281/zenodo.10210210. Pooled-sequencing reads are available at NCBI under project number PRJNA1045841.

## Acknowledgements

Funding was provided by the Swiss National Science Foundation (Grant No: IZCOZO_177062). We thank the Functional Genomic Centre Zurich (FGCZ) for their support in library preparation and sequencing. We are grateful to Christoph Grieder and the Fodder Crop Breeding group of Agroscope Zurich-Reckenholz for infrastructure and support with the generation of the F_2_ population and the *Xtg* resistance screening.

## Author contribution

Study design: FG, VK, RK, BS. Data collection and analysis: FG, with the help of YC, SY, DC. Writing and reviewing: FG, RK, BS. All authors contributed to the article and approved the submitted version.

## Supplementary information

**Supplementary figure 1** Association of each variant locus to *Xanthomonas translucens* pv. *graminis* resistance in *Lolium multiflorum* calculated with the t-|ΔSNP| index (A, B), the G’ value (C, D) and the ED^4^ value (E, F), with Rabiosa as a reference genome sequence, on all pseudo-chromosomes (A, C, E) or the region with the highest association to resistance on pseudo-chromosome 4 (B, D, F).

Dotted blue lines represent the 95% confidence interval for the t-|ΔSNP| index, and dotted red lines represent the 99% confidence interval for the t-|ΔSNP| index or the false discovery rate threshold of 1% for the G’ value.

**Supplementary Table 1** Genes found within the candidate region, defined from 28,800,000 bp to 28,900,000 bp on pseudo-chromosome 4 of the M2289_v2 genome sequence, and from 163,090,000 bp to 163,410,000 bp in the Rabiosa genome sequence, that could not be annotated or could not be linked to disease resistance.

---

**Supplementary Figure 1:**
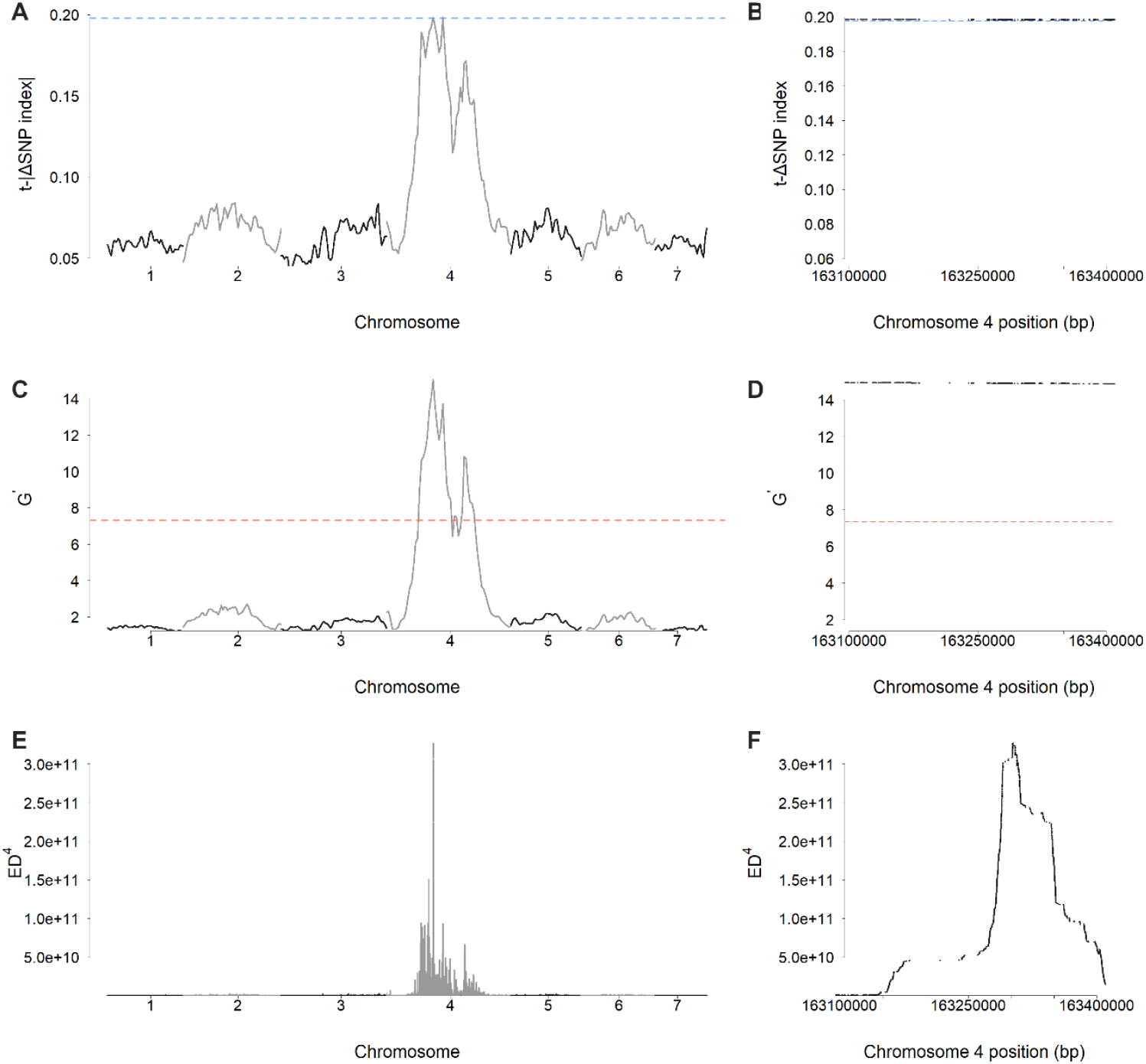

**Supplementary Table 1:**
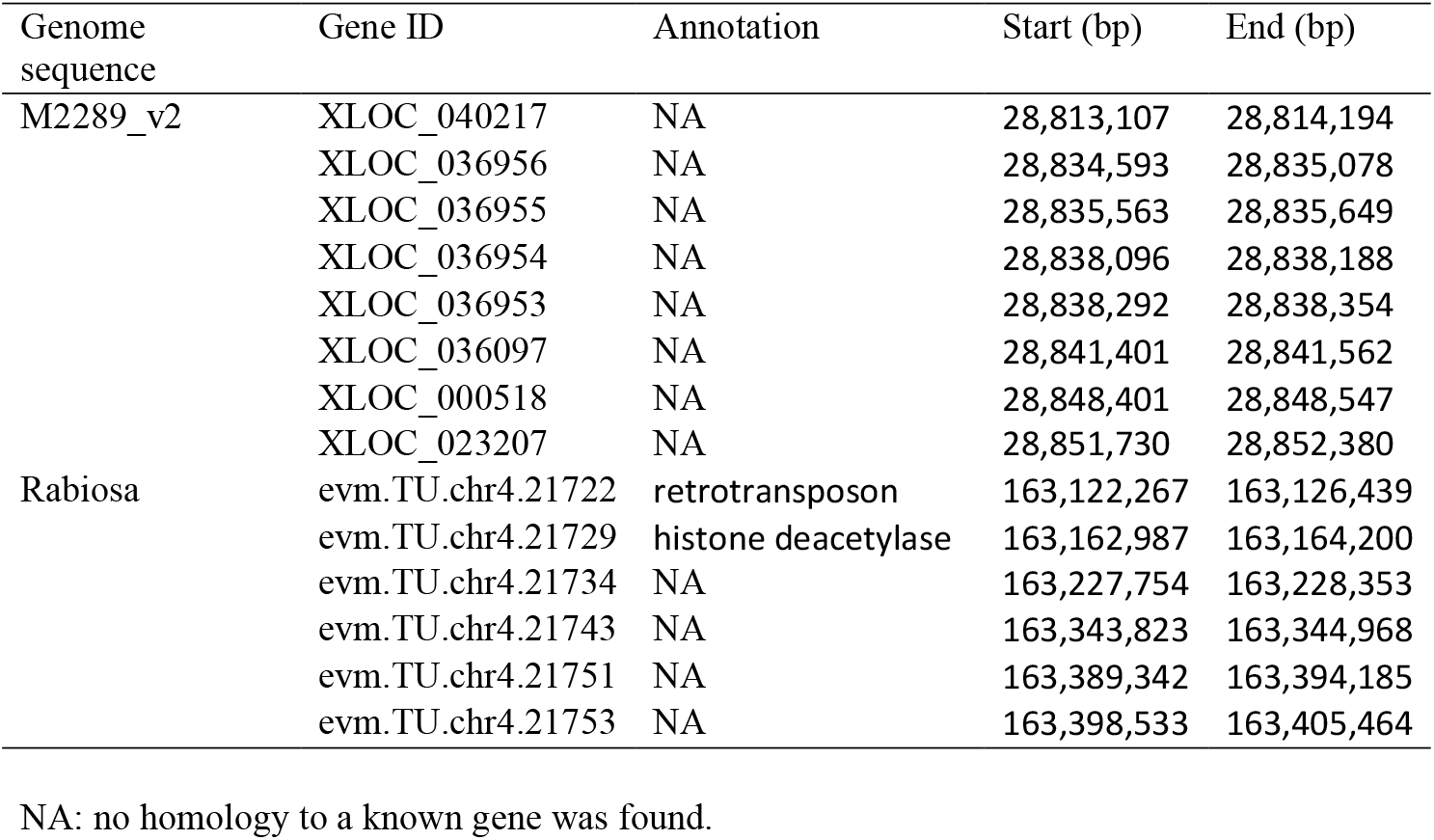

| Genome<br>sequence | Gene ID | Annotation | Start (bp) | End (bp) |
| --- | --- | --- | --- | --- |
| M2289_v2 | XLOC_040217 | NA | 28,813,107 | 28,814,194 |
|  | XLOC_036956 | NA | 28,834,593 | 28,835,078 |
|  | XLOC_036955 | NA | 28,835,563 | 28,835,649 |
|  | XLOC_036954 | NA | 28,838,096 | 28,838,188 |
|  | XLOC_036953 | NA | 28,838,292 | 28,838,354 |
|  | XLOC_036097 | NA | 28,841,401 | 28,841,562 |
|  | XLOC_000518 | NA | 28,848,401 | 28,848,547 |
|  | XLOC_023207 | NA | 28,851,730 | 28,852,380 |
| Rabiosa | evm.TU.chr4.21722 | retrotransposon | 163,122,267 | 163,126,439 |
|  | evm.TU.chr4.21729 | histone deacetylase | 163,162,987 | 163,164,200 |
|  | evm.TU.chr4.21734 | NA | 163,227,754 | 163,228,353 |
|  | evm.TU.chr4.21743 | NA | 163,343,823 | 163,344,968 |
|  | evm.TU.chr4.21751 | NA | 163,389,342 | 163,394,185 |
|  | evm.TU.chr4.21753 | NA | 163,398,533 | 163,405,464 |

NA: no homology to a known gene was found.

